# Drought reduces solitary bee reproduction and skews sex ratios

**DOI:** 10.64898/2026.05.29.728802

**Authors:** Lindsie M. McCabe, Byron Love, Jonathan B. Uhuad Koch, Kelsey Graham, Diana Cox-Foster

## Abstract

The Intermountain West of the United States has experienced years of extreme drought and increased temperatures. Increasing temperature and droughts can negatively impact native species that are locally adapted to environmental conditions that have persisted prior to the Anthropocene. Bees, especially solitary bees, provide critical ecosystem services because they pollinate ~90% of all flowering plants. Here we asked how drought impacted the reproduction of *Osmia bruneri*, a solitary mason bee. We released 20 females in nesting blocks at 21 field sites across four years (2020 – 2023) in the Bear River Mountains of northern Utah. *O. bruneri* reproduction was positively correlated with winter precipitation, and sex ratio was skewed in years that had a yearlong drought from the expected female to male ratio of 1.3:1 to 0.33:1. These results suggest that *O. bruneri* is susceptible to winter precipitation droughts. Not only was there a decrease in the total number of cells provisioned but the overall number of females produced decreased significantly during season long drought conditions. Continuous droughts can lead to a local level population decline and could contribute to overall species declines. Identifying the effects of extreme drought on solitary bee fecundity is critical for supporting effective management practices and conservation prioritization. Additionally, the results suggest that preceding winter precipitation can act as an indicator for predicting nesting success in wild solitary bees and may be an overall indicator of habitat quality.

## Introduction

Increasing temperatures and decreased precipitation, as well as variable weather conditions has emerged as a critical concern across the globe, with far reaching impacts on ecosystem services and biodiversity, including pollinators and the plants they depend upon (Muluneh 2021, Outhwaite et al 2022). These extreme abiotic events as well as changes in plant phenology have been especially disruptive in the western United States, which is currently experiencing a multi-decade drought (Biederman 2024). Plant communities in the montane and arid ecosystems of this region store precipitation in the form of winter snow, released during spring and early summer runoff (Fritze et al., 2011). As a result, the timing of snowmelt is of equal importance to annual snowpack as drivers of seedling recruitment, plant growth and reproduction (Comstock & Ehleringer 1992), which in turn have cascading effects on diversity and ecosystem function (Sun et al 2015, Kudo 1992, Jerome et al 2021). For example, water scarcity can stunt plant growth, reduce pollen quality and nectar concentrations and influence gene expression (Davies et al 1986, Bhat et al 2024, Ahluwalia et al 2021), all of which are factors that have been shown to affect bee foraging strategies (Rering et al. 2020, Kuppler et al. 2021). Likewise, drought can lead to reduced reproduction and fitness in bees, which have strong biotic ties to their host plants, relying on them for both pollen and nectar resources. Declines in bee diversity in the year or years following drought are well documented (Thomson 2016, Philips et al 2018, Hung et al 2021, LeBuhn and Luna 2021).

Bees are keystone species, providing up to 90% of all pollination services (Potts et al, 2010; 2016, Ollerton et al. 2011). Changes in local bee community dynamics (i.e., bee abundance, species richness and phenology) can therefore be used as an indicator of ecosystem health (Goulson, and Nicholls 2016). Most bee species in temperate regions are univoltine, with a single generation produced each year. Consequently, the adult bee community each year is the direct result of conditions experienced the year before. For example, Kammerer et al (2021) found that wild bee abundance and richness were best predicted by the previous year’s weather, and Pardee et al. (2022) reported bee traits such as body size, diet breath, nesting substrate and overwintering stage were all strongly correlated with current and previous climate variables (mean annual temperature, winter precipitation, and mean annual precipitation). This rapid response underscores the importance of understanding how climate variables affect bees, which in turn gives us stronger predictive capabilities on changes in local bee communities.

*Osmia bruneri* Cockerell, 1897 (Hymenoptera: Megachilidae) is a solitary cavity nesting bee active during late spring through mid-summer, unlike many other *Osmia* which are mostly active in the early spring (Frohlich 1983). They are known to nest in pre-existing cavities in dead wood and plant stems and will readily use artificial nest substrates such as straw inserts placed in drilled wood blocks (Frohlich 1983). This species is generally thought to be a generalist pollinator but with preferences towards *Phacelia* (Boraginaceae) (Cripps and Rust, 1989). *O. bruneri* makes a good model system to answer questions about the effects of climate and drought due to their ability to nest in cavities, a heightened flight period during the summer months (usually June – August), and a generalist pollen diet. We predicted that *O. bruneri* would decrease nesting in years of drought, and that sex ratio would be skewed towards males when conditions were unfavorable (i.e. lack of precipitation). Many bee species have shown the sex ratio can be influenced by the environmental conditions around them. In particular *O. bruneri* sex ratio is known to change based on the season, In the early summer, *O. bruneri* can produce more females than males, but late in the summer they produce more males than females (Frohlich and Tepedino 1986). This could suggest that later in the summer when resources become scarce that resources limitation is driving the sex ratio difference, supporting sex allocation theory and bet-hedging strategies in solitary bees (Danforth 1999). However, Peterson and Roitberg (2006) found that *Megachile rotundata* foraging in limited resources areas allocate more resources to females but not males, which lead to a skew female bias sex ratio and underdeveloped males. Which demonstrates more investment placed into females first and then males. The results of our study can serve as a framework in demonstrating the role of drought on sex ratios in solitary bees and have implications for managing solitary bee pollinators on wildlands

## Methods

Study Site: The Bear River Mountains are a subset of the Wasatch Mountains (which in turn are a subset of the Rocky Mountains) straddling the northeastern Utah-Idaho border. Made primarily of limestone and dolomite, elevations range from 1,425 – 3,042 m asl. The climate is typical of semi-arid, high-desert mountains of the Intermountain West, with most precipitation coming from winter snowfall (avg 50 – 75 mm each winter), which melt throughout spring and into early summer (See table 1). Plant communities include sagebrush-steppe in flat valleys and southern exposures, and aspen-conifer forests on northern exposures and higher elevations.

**Table 1:** Tabel showing site locations, location of the canyon and the elevation at each site. Mean annual precipitation is recorded for all four years plus a 30-year norm. Blank spaces in the year column indicate that that site was not used that year. Colors within the MAP columns indicate levels of precipitation (Red = below average precipitation, Yellow = average precipitation, and Green = above average precipitation).

| Site | Canyon | Elevation (m) | MAP 2020 (cm) | MAP 2021 (cm) | MAP 2022 (cm) | MAP 2023 (cm) | 30-year MAP (cm) |
| --- | --- | --- | --- | --- | --- | --- | --- |
| BF1 | BF | 2237 | 51.33 | 43.33 | 56.12 |  | 54.43 |
| BF2 | BF | 2215 | 51.32 | 43.22 | 56.07 |  | 54.38 |
| BF3 | BF | 2191 | 51.50 | 43.26 | 56.16 |  | 54.47 |
| BF4 | BF | 2148 | 51.58 | 43.08 | 55.56 |  | 53.89 |
| BF5 | BF | 2197 | 51.52 | 42.70 | 54.43 |  | 52.79 |
| BF6 | BF | 2206 |  | 45.33 | 63.76 |  | 61.84 |
| BF7 | BF | 2218 |  | 43.67 | 55.19 |  | 53.53 |
| FB1 | FB | 2312 |  | 53.94 | 69.49 | 100.61 | 67.40 |
| FB2 | FB | 2306 |  | 53.41 | 68.92 | 102.45 | 66.85 |
| FB3 | FB | 2321 |  | 54.50 | 70.11 | 114.13 | 68.01 |
| FB4 | FB | 2470 |  | 65.51 | 82.14 | 121.25 | 79.67 |
| FB5 | FB | 2487 |  | 66.43 | 83.24 | 124.24 | 80.74 |
| FB6 | FB | 2511 |  | 67.63 | 84.76 | 124.50 | 82.21 |
| FB7 | FB | 2515 |  | 67.86 | 85.04 | 125.23 | 82.48 |
| TC1 | TC | 2090 |  | 43.75 | 56.80 | 81.79 | 55.09 |
| TC2 | TC | 2107 |  | 44.23 | 57.31 |  | 55.97 |
| TC3 | TC | 2091 |  | 45.00 | 58.59 |  | 56.83 |
| TC4 | TC | 1955 |  | 39.81 | 51.19 | 76.27 | 49.65 |
| TC5 | TC | 1945 |  | 38.85 | 49.66 | 73.49 | 48.17 |
| TC6 | TC | 1980 |  | 39.54 | 50.31 | 81.34 | 48.87 |
| TC7 | TC | 2029 |  | 40.51 | 51.20 |  | 49.36 |

A total of 21 sites were selected in three canyons across the northern Utah portion of the range on wildlands managed by the Wasatch-Cache US Forest Service, Blacksmith Fork, Twin Creek, and Franklin Basin (SI1 & Table 1). Sites ranged in elevation from 1,793 – 2,325m. In 2020 (year 1) n=5 sites were sampled in Blacksmith Fork canyon. In 2021 and 2022, (years 2 and 3) n=21 sites were sampled from all three canyons. In 2023 (year 4), n=11 sites were sampled from Franklin Basin and Twin Creeks (SI1). While native wildlife and plants are distributed across the study area, the U.S. Forest Service units are public lands and thereby mandated to support grazing habitat for livestock and outdoor recreational activities. The site sampling schedules were determined by funding resources and personnel availability. Sites were sampled every two weeks starting roughly two weeks after the last snow melt (early to mid-June) through the end of August.

Study design: Nest blocks, an 8 × 8 × 15 cm wood block with 16-4mm diameter holes with paper straw inserts, were placed at each site on metal fenceposts approximately 1m high. Nests were stocked with four *O. bruneri* nests (straws) containing a total of 20 females and 20 males. Prior to release, nests were x-rayed to determine the health of the bees (i.e. bees had adequate fat bodies, were adults, and were not parasitized). *O. bruneri* stocked nests along with nest block were placed out each year according to snow melt and bloom time for each year: year 1 = 6 June 2020, year 2 = 20 June 2021, year 3 = 17 June 2022, and year 4 = 28 June 2023 (SI.1). While we do not know for certain if all newly made nest were from our released bees or *O. bruneri* naturally in the system, we were overall interested in the amounts of nests made and cells produced per nest, not estimating total populations for each site. Furthermore, *O. bruneri* is known to be philopatric and not disperse from where their natal nests, so it is more likely these nests were made by bees we released (Frohlich 1983). Nest blocks were inspected every other week, and completed nests were replaced with empty straws. Completed nests were pulled from the nest block if the outside of the straw was capped with masticated leaves. All completed non-*O. bruneri* nests were also removed to not induce nesting competition, given that other bee and wasp species also used these nesting cavities. Completed straws were brought back to the USDA-ARS Pollinating Insects Research Unit (PIRU) in Logan, UT to complete development under local ambient temperature (23 – 32° C). A radiograph of each nest was developed to confirm the number of cells provisioned and monitor development. Furthermore, a radiograph was achieved for all completed nests upon being brought in to confirm the bee species, the number of cells made, and the contents of the cells. We used a Faxitron 48804N, (Faxitron Bioptics, Tuscon, AZ) to capture a radiograph of each bee nest at 28 KV for 15 seconds. Unfinished nests or abandoned nests (i.e. not capped) were left out for the duration of the season and were analyzed at the end of the season with a radiograph.

On November 1^st^ of each year, all nests were placed into storage for the winter at 4°C. Cells remained in their original nests through the duration of winter. In the late spring/early summer overwintering cells were stripped from their nests and placed into gel capsules to monitor emergence and confirm species identifications and sex. Cocoons were then placed into incubators to replicate ambient early summer temperature to induce emergence. Only nests made by *O. bruneri* were considered for this study, nests made by other *Osmia* species or other bee/wasps were not assessed.

In addition to the nest blocks being checked every two weeks, floral evaluations were also monitored. Three belt transects consisting of 1 × 30 m were walked every two weeks to document plant species and flower abundance for each site for the duration of the project. Plants were identified by the authors and confirmed by a local forest service botanist, Patricia Winn, North Zone Botanist for Uinta-Wasatch-Cache National Forest. Transects were within 50 yards of each nest block.

Climate Data: Temperature and precipitation data were extracted from the PRISM Group, which gathers data from monitoring networks to reveal long and short term weather patterns (https://prism.oregonstate.edu), for each of the sites and years at an 800 m resolution. Temperature data were validated with on-site/on-nest iButton data logger (iButtonLink). Snow water equivalent (SWE) data was taken for each of the canyons using SNOTEL (Snow Telemetry), a series of systems throughout the western United States used to measure snowpack and precipitation (https://wcc.sc.egov.usda.gov/nwcc/). Data from three SNOTEL sites, Bug Lake, Temple Fork, & Klondike Narrow, which where the closest SNOTEL site to the sites that we samples. All sites in each canyon were within 3 km from their SNOTEL sites. All weather data (temperature, precipitation, and SWE) was analyzed based on “mean annual” data with the start of the year being October instead of January, so our climate data were 2020: Oct 2019 – Sep 2020, 2021: Oct 2020 – Sept 2021, 2022: Oct 2021 – Sept 2022, and 2023: Oct 2022 – Sept 2023. Our aim for this data partitioning method is to better account for fluctuations of winter precipitation on floral resources and reproduction of *O. bruneri*.

Statistical Analysis: Because of the strong biotic and abiotic association that go into bee reproduction, we constructed two piecewise structural equation models (SEMs) to test for an effect between abiotic effects (precipitation, temperature and elevation) and floral resources (plant abundance and plant richness) on *O. bruneri* reproduction (cells provisioned and nest provisioned). The SEMs allowed us to analyze the complex relationships between weather and floral resources and test for both direct and indirect effects not available in traditional regression analyses. Cell provisioning was defined as the number of cells laid in each nest and nest provisioning was defined as the number of nests made at each site. Initial hypothesis models included winter precipitation, maximum SWE, summer precipitation, mean annual precipitation, mean annual temperatures, floral abundance, floral richness and number of cells provisioned or number of nests provisioned (Fig 1). The final model included only winter precipitation, floral abundance, and number of nests provisioned. Additionally, we constructed a generalized linear mixed model to test if sex ratios differed between years and included canyon as a random effect. All data were analyzed in R 4.0.1 using packages piecewiseSEM (Lefcheck et al 2021), glmm (Knudson 2022), and ggplot2 (Wickham et al 2016).

**Figure 1:**
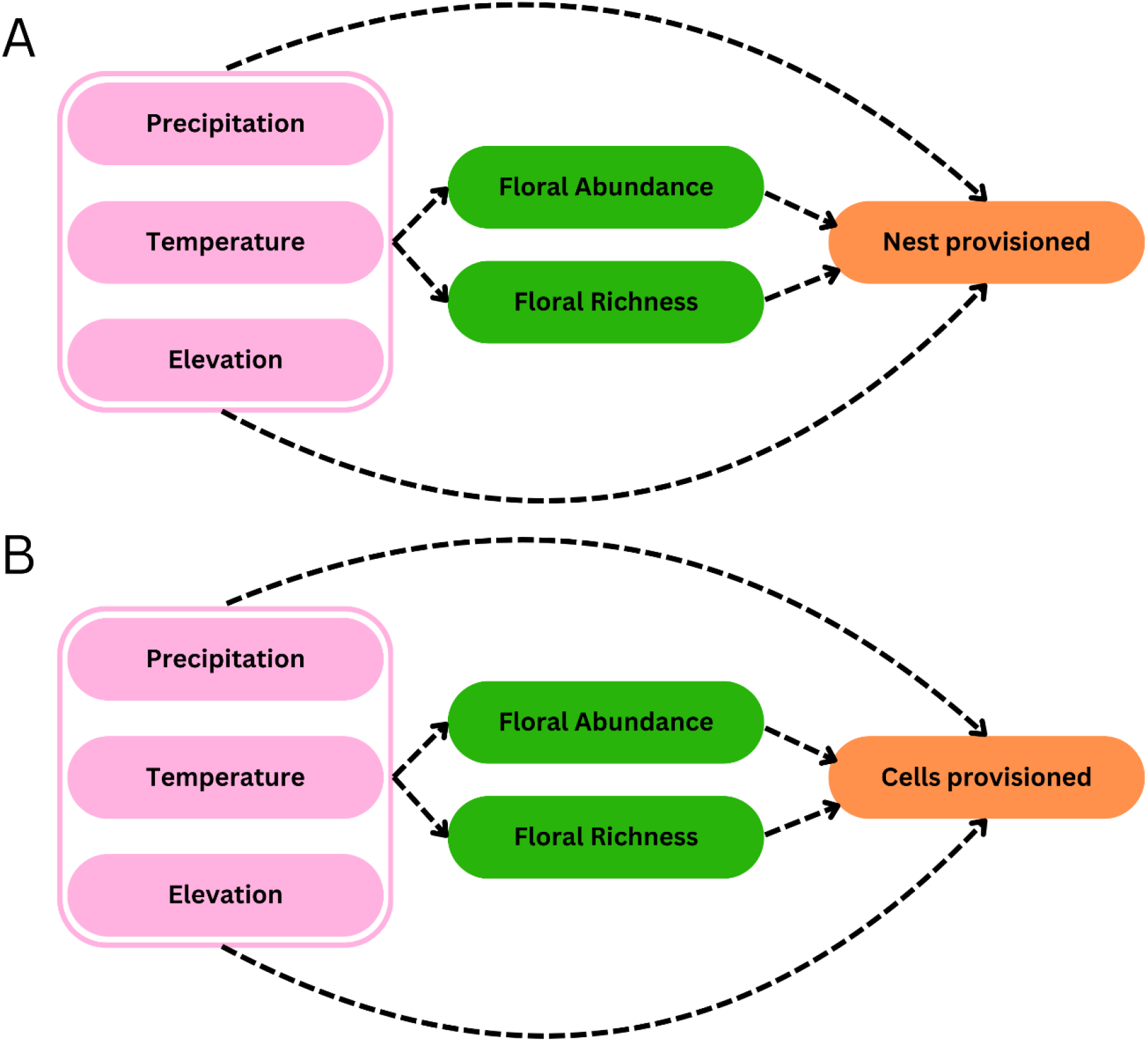
Concept model for pathway interactions. Precipitation variables including mean annual precipitation, winter precipitation, summer precipitation, and maximum snow water equivalent.

**Figure 2:**
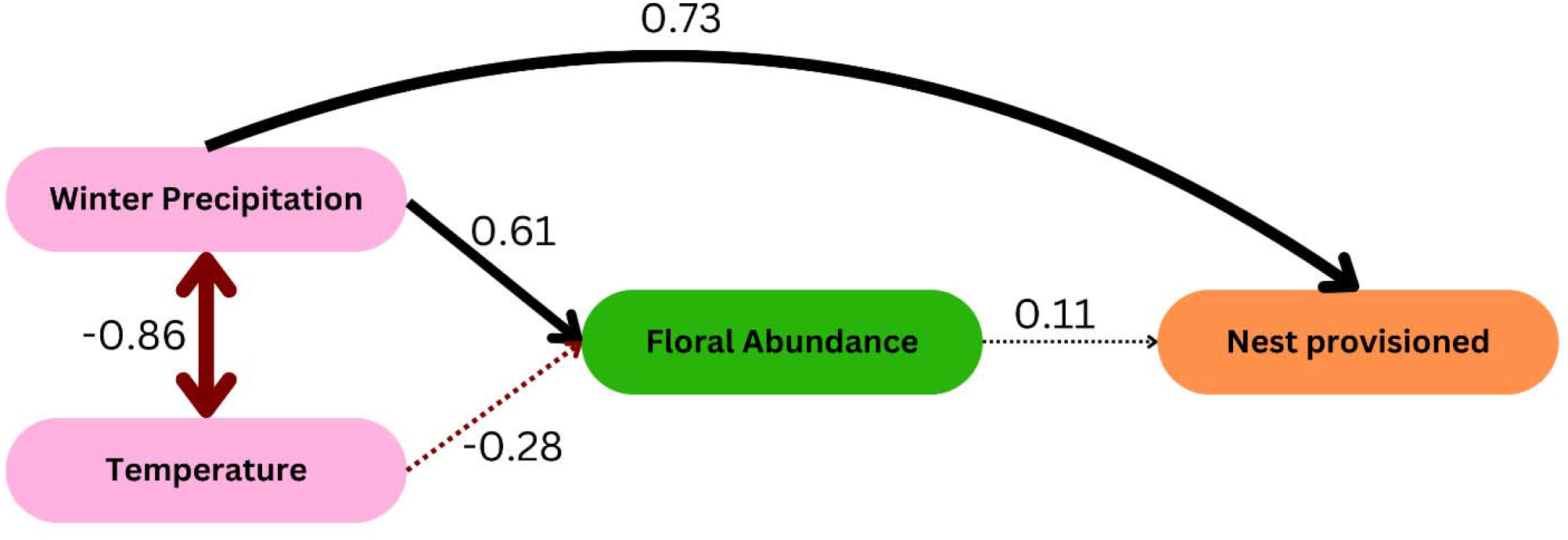
Pairwise structural equation model showing pathways of interactions. Width of the bar and numerical number represents R^2^ values, where solid lines are significant and dashed lines are non-significant interactions. Arrows denote directionality, red arrow signify negative relationships, while black arrow signify positive relationships.

## Results

A total of 609 nests with 2,859 cells were collected in our study for 2020-2023. For completed nests, nests had 1 to 14 cells with an average of 4.01 ± 1.9 SD cells per nest. Additionally, we tracked a total of 117 plant species over the course of the four years. The best-fit model (Fisher’s C = 31.56, p = 0.956, AICc = 812) had no missing paths and had a substantially better fit than our full model with all hypothesized links between variables (AICc = 1644). The SEM model that had the best AIC included winter precipitation (over mean annual, summer precipitation, or maximum SWE), floral abundance (but not floral richness) and mean annual temperature. We found that increased winter precipitation had the greatest impact in positively influencing bee reproduction (number of completed nests and number of cells produced) (unstandardized coefficient = 2.145, p = 0.016; Fig.2). The model predicts that for each one cm increase in winter precipitation, we would expect 2.14 more cells produced. Increased winter precipitation was correlated with increased plant abundance (unstandardized coefficient = 2.390, p = 0.017); but surprisingly, increased plant abundance did not have a direct effect on bee reproduction. Floral abundance was retained in the model but had a non-significant effect (unstandardized coefficient = 0.423, SE= 0.004, p = 0.673).

In 2021 the Intermountain region experienced the worst drought since 2012 (SI.2). When mean annual precipitation (MAP) was below the 30-year average, below-average MAP had a negative effect on the sex ratio of *O. bruneri* (Figure 3). In years at or above MAP, the sex ratio was 1.33 females for every one male produced; however, in years under the MAP (i.e., 2021) the sex ratio was reduced significantly by ~79% with one female for every three males produced (z = 44.08, p < 0.001).

**Figure 3:**
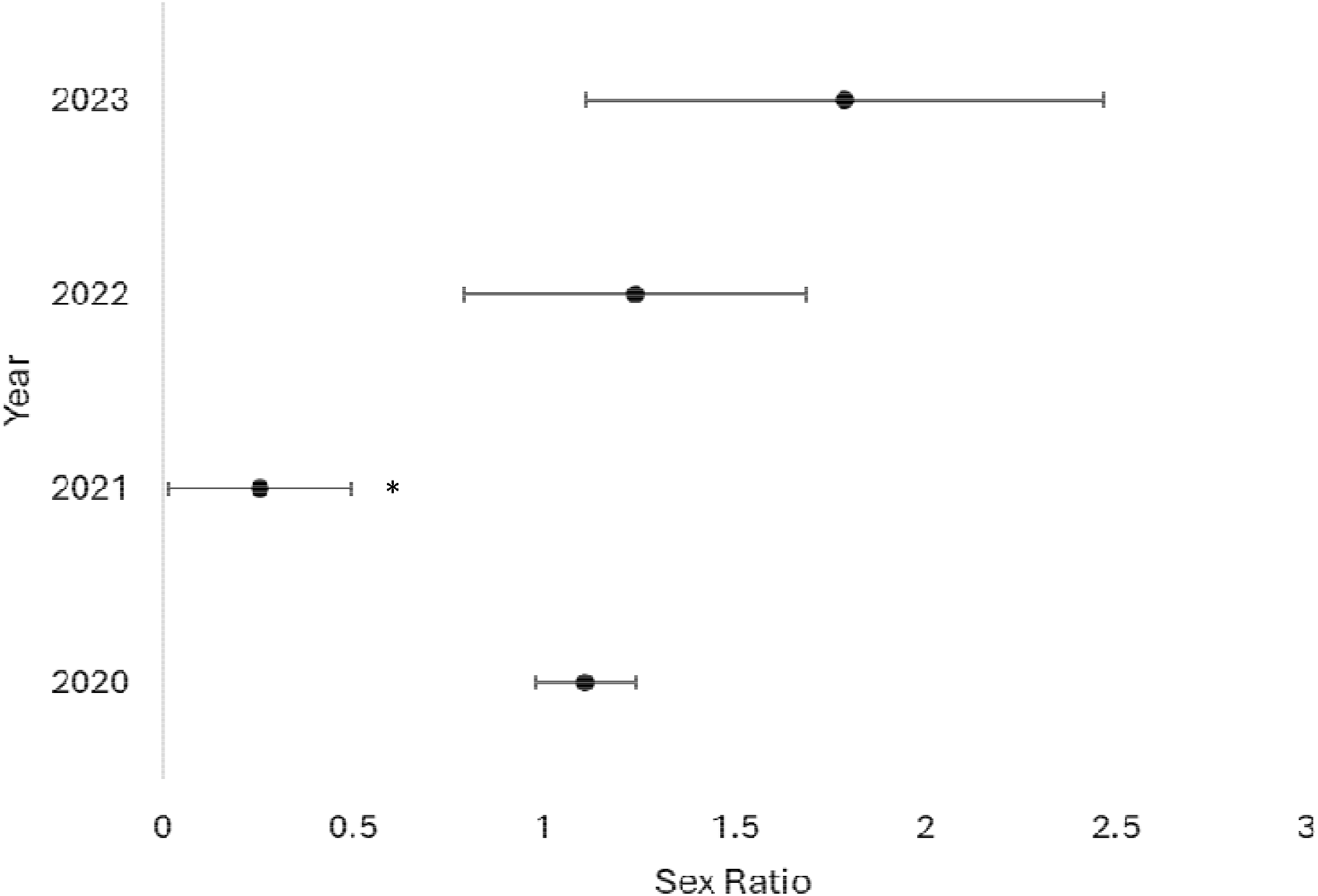
Forest plot showing the average number of females produced for every one male during each year of the study. Bars represent standard error. Asterisk denotes significant difference.

## Discussion

Our results demonstrate that *O. bruneri* fecundity is susceptible to changes in winter precipitation and that *O. bruneri* may be used as indicator species for environmental health of the ecosystem. Extreme drought amplifies reproductive loss, resulting in significantly fewer individuals being produced and, in a male-biased sex ratio in the offspring. As droughts are likely to become more frequent, (Strzepek et al 2010, Xu et al 2019), droughts have serious implications for the health and longevity of solitary bee populations, especially in arid regions like the Intermountain West of the U.S. Furthermore, because we directly measured reproductive output, solitary bees can be used as a model system to predict their populations in the years to come.

Previous studies (Kazenel et al 2024, Pardee et al. 2022, Kammerer et al 2021, Soroye et al 2020) attempted to demonstrate reproductive output by estimating adult bee abundance from pan traps or sweep netting and using previous years weather variables. However, by using wild cavity nesting bees deployed in nest blocks, we directly measured reproduction in response to ambient environmental stressors. Moreover, reproductive success is a measure of fitness and therefore can be applied to population models and simulations. The data generated in our study is novel, demonstrating not just which species and individuals survive to adults and forage the following year but directly measuring reproductive effort and sex ratio of offspring.

Surprisingly, we found no significant relationship between plant abundance and reproduction, but there was a strong positive relationship between plant abundance and winter precipitation. We hypothesize that *O. bruneri* are unlikely to use all or even most floral hosts represented in our plant survey, which may have limited our ability to capture direct impacts of reduced floral availability on reproductive output. Drought and temperature may also affect the physiology of the plants and the nutritional quality of their pollen, this can look like reduced flowers produced, reduced pollen or nectar availability or altered flowering times (Buchmann and Papaj 2024, Brunet et al 2024). Downstream effects of these traits may impact bee reproduction and provisioning and later impact the bee’s physiology and the egg quality.

Additionally, there may have been other resource constraints, such as nesting resources, that were driving reductions in reproductive output. *O. bruneri* use leaf pieces and then masticate them to form their nest cells (Cane et al 2007). In times of increased drought, plants often harden their leaves, and this could cause preferred plants to be unusable (Nardini 2022). Unfortunately, little is known about what plants these bees used for nest construction nor about the overlap between biotic nesting and food resources. Future research is needed to identify preferred host plants for pollen and leaves and how changes in their availability impact *O. bruneri* reproductive effort.

Initial work on sex ratio of *O. bruneri* suggested an average 1:1.35 female skewed sex ratio was normal for *O. bruneri* (Frolich 1986). Our work found that in years with above average mean annual precipitation that the sex ratio average was 1:1.6 with no significant differences between years, except in 2021 where sex ratio was 1:0.33 with the drought. Given our results, we hypothesize that the optimal sex ratio of *O. bruneri* population is at an ecosystem equilibrium (i.e. more precipitation does not mean more females laid versus males). However, in a stressed environment it is likely that mothers are choosing to provision and lay male cells over female cells since their resources needs are less. This has been seen in other groups such as Megachile rotundata which exhibited high levels of males being laid when resources were limited (Peterson et al 2006); although, there is a correlation between adult fitness of the mother and the number of female offspring she produces (Peterson 2006). The change in sex ratio with changes in resources or environmental conditions may be an example of bet hedging in uncertain environments (Danforth 1999). Whereby the mothers are choosing to sacrifice reproductive females, in hopes that at least the males will survive. Bet-hedging can be a strategy used by species in highly variable environments to increase the risk of survival (Hairston and Fox 2013). Bet-hedging has been previously found in bees as a strategy to avoid long periods of dry conditions (Danforth 1999). However, bet hedging usually is exhibited in the individual’s ability to be plastic during diapausing periods (Chesson 2000) and not a switch in sex ratio. Our results point to the need to understand how community and population dynamic changes over time and through multiple generations under adverse weather conditions.

Similar drought-related declines have been observed in other *Osmia* species using trap-nest methods (e.g., Bosch & Kemp 2000; Peterson et al. 2006). For example, arid conditions were found to limit nest occupation and brood productivity in *Osmia cornuta* (Bosch et al. 2023). If other solitary bee species follow similar patterns as observed in our study, there could be substantial impacts on the overall bee communities in years of drought and cascading effects on the environment. Reduced and sex-skewed reproductive output may have negative impacts on the ecosystem services that bees provide to plants in both natural and agricultural ecosystems. *Osmia* is an abundant genus of solitary bees in montane environments of the Intermountain West. Thus, identifying how winter droughts and other environmental stressors effect these cavity nesting bees is critical for supporting ecosystem services. Furthermore, reduced fecundity and male-biased sex ratios could be a limitation for wild propagation efforts for cavity nesting bees used in commercial pollination, like *Osmia lignaria* (Bosh and Kemp 2000, Pitts-Singer et al 2018, McCabe et al 2024). As such, sex ratio of offspring in nests could be used as a metric for commercial operations to determine overall health of wild and managed populations of *Osmia*. Decreased female *O. bruneri* reproduction can lead to reduced genetic diversity in this population which could further the risk of local extinction (Frankham 2005). Reduced genetic diversity has been linked to local extirpation in plants such as the *Clarkia pulchella* (Newman and Pilson, 1997) as well as insects, such as the Glanville fritillary butterfly (Saccheri et al 1998). Additionally, increased inbreeding in naturally outbred species leads to inbreeding depression (Ralls et al 1988). Reduced and skewed reproductive output can reduce genetic diversity in a population, which we demonstrate is exacerbated by drought conditions in the western United States.

Solitary cavity-nesting bees experience climate and weather events differently than solitary ground nesting bees or social bee species due to differences in behavior and life history traits. Ground nesting bees, including overwintering bumble bee gynes, are typically buffered from cold winters because of how deeply they are buried in the ground (Vaughan and Black 2008). On the other hand, social bee colonies use the overall numbers in the colony to regulate responses to weather events, such as warming temperatures (Ostwald et al 2022, Davison and Fields 2018).

We propose that winter precipitation can act as a carrying capacity indicator for solitary cavity-nesting bees for the upcoming season, meaning that more winter precipitation can lead to more bee reproduction in the spring and summer. Because of the predictable influence of winter precipitation on solitary bee fecundity, winter precipitation levels may provide solitary bee producers and land managers with a way to predict levels of propagation of native *Osmia* spp. in the Intermountain West. However, potential management decisions would benefit from research replicating our study design across the Intermountain West to ascertain the impacts of winter precipitation on solitary bee fecundity. Fortunately, propagating bees, particularly *Osmia* spp. with nest blocks can be highly successful, and in the case of native *O. bruneri* replicable in montane environments across multiple years.

---

**Supplemental Figure 1:**
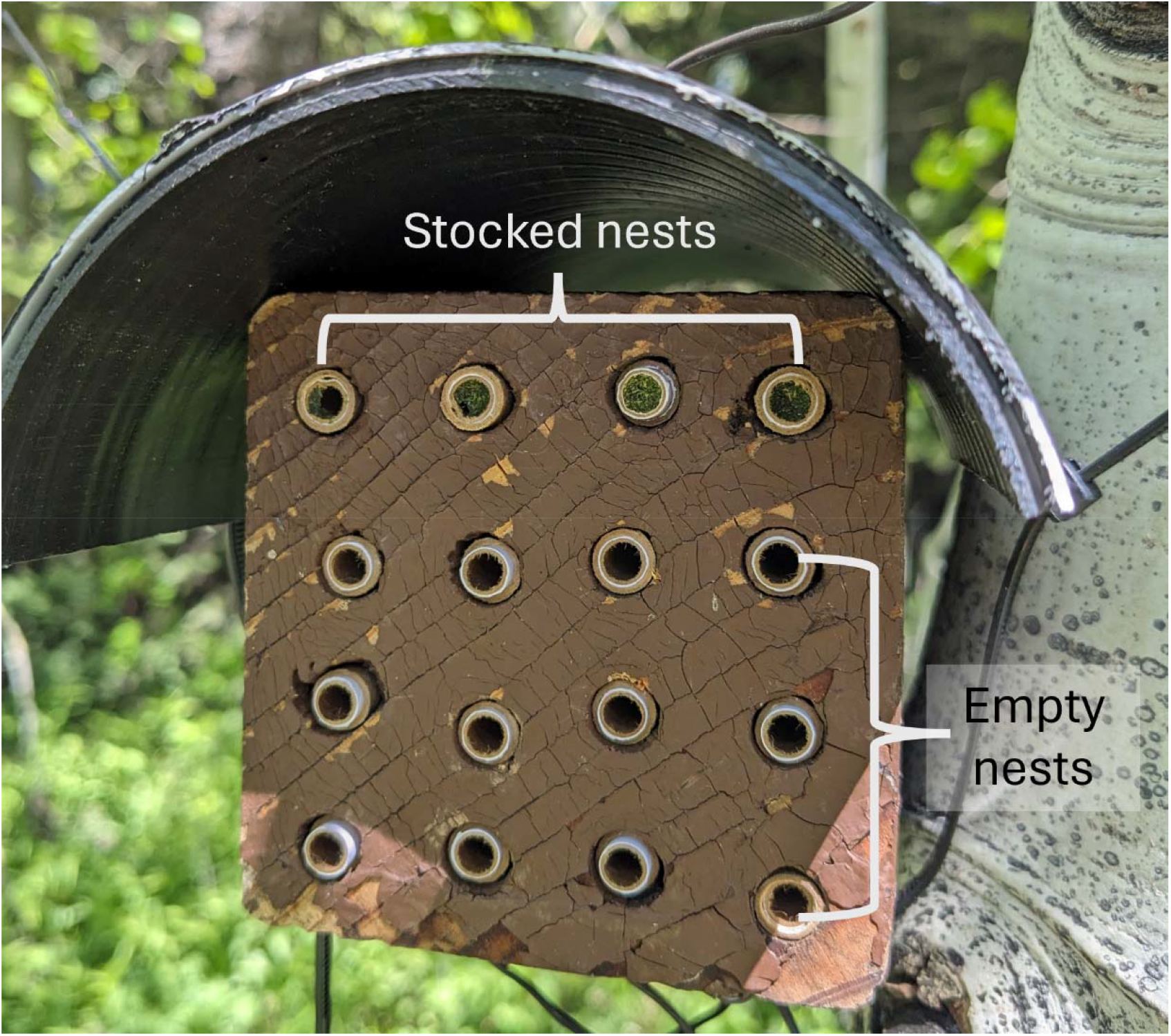
Example nest block that was put up at each site. Top row of each block had four stocked nested with approximately 20 bees per block (distribution not always equal between straws). These top straws were replaced with unused straws after emergence (usually after the first week). The bottom 12 straws were empty and ready for nesting.

**Supplemental Figure 2:**
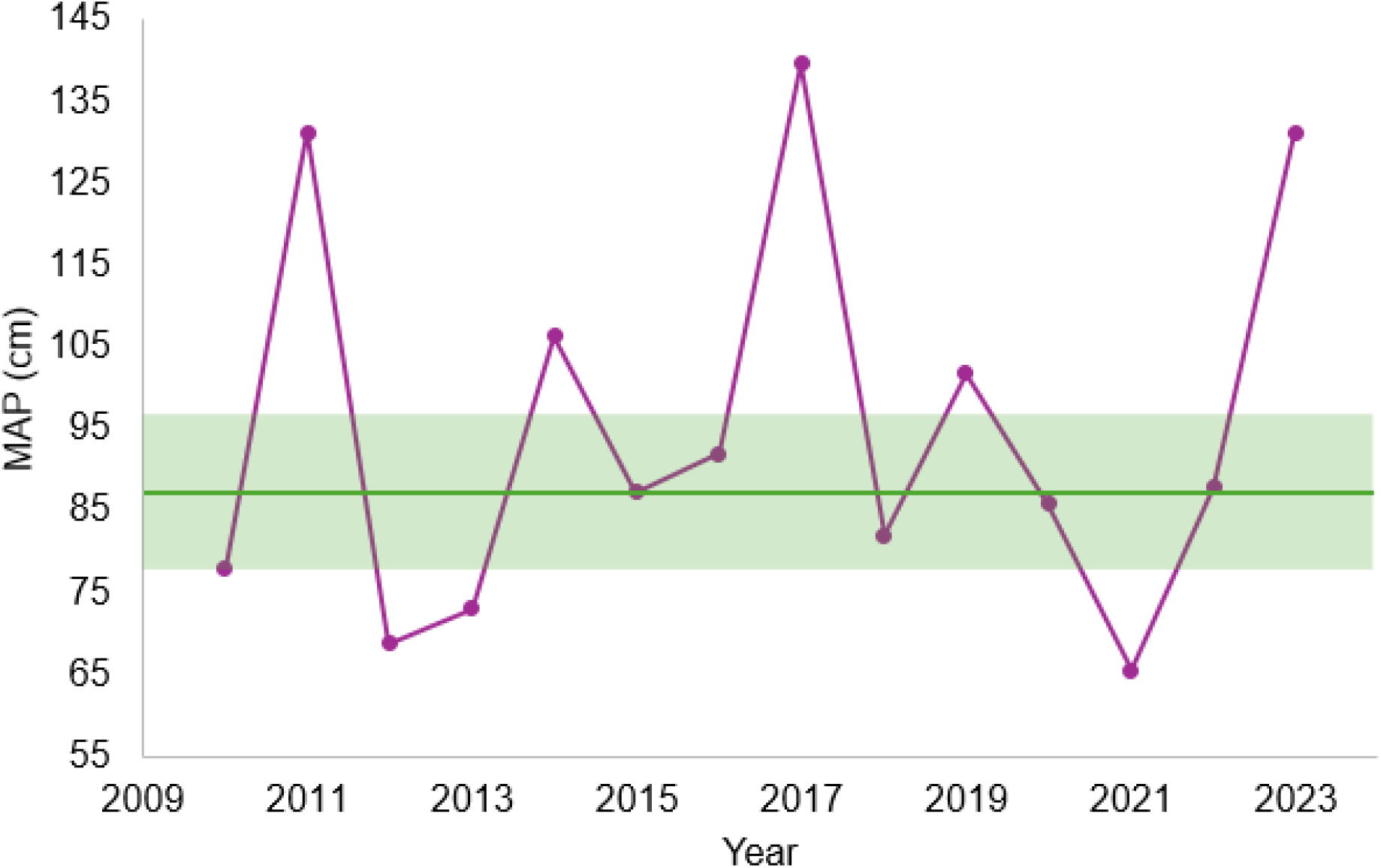
Mean annual precipitation for Logan Canyon from 2010 – 2024. Data extracted from the USDA NRCS Climate and Hydrological Norms data set. Dark green bar indicated the rolling average MAP from 1991 – 2020 and the light green shaded areas represent the two standard deviations around the mean.

## Notes

### Competing Interest Statement

The authors have declared no competing interest.

### Funding

- Project Apis m.

## Literature Cited

Ahluwalia, O., Singh, P.C., Bhatia, R. (2021). A review on drought stress in plants: Implications, mitigation and the role of plant growth promoting rhizobacteria. Resources, Environment and Sustainability, 5, 100032.

Biederman, J.A., Zhang, F., Dannenberg, M.P., Yan, D., Reed, S.C., Smith, W.K. (2024). Reply to Comment on “Five decades of observed daily precipitation reveal longer and more variable drought events across much of the western United States”. Geophysical Research Letters, 51(1), e2023GL105124.

Bhat, M.A., Mishra, A.K., Shah, S.N., Bhat, M.A., Jan, S., Rahman, S., Baek, K.-H., Jan, A.T. (2024). Soil and Mineral Nutrients in Plant Health: A Prospective Study of Iron and Phosphorus in the Growth and Development of Plants. Current Issues in Molecular Biology, 46(6), 5194–5222.

Bosch, J., Kemp, W.P. (2000). Development and emergence of the orchard pollinator *Osmia lignaria* (Hymenoptera: Megachilidae). Environmental Entomology, 29(1), 8–13.

Cane, J.H., Griswold, T., Parker, F.D. (2007). Substrates and materials used for nesting by North American *Osmia* bees (Hymenoptera: Apiformes: Megachilidae). Annals of the Entomological Society of America, 100(3), 350–358.

Chesson, P. (2000). Mechanisms of maintenance of species diversity. Annual Review of Ecology and Systematics, 31(1), 343–366.

Comstock, J.P., Ehleringer, J.R. (1992). Plant adaptation in the Great Basin and Colorado plateau. The Great Basin Naturalist, 52(3), 195–215.

Cripps, C., Rust, R.W. (1989). Pollen foraging in a community of *Osmia* bees (Hymenoptera: Megachilidae). Environmental Entomology, 18(4), 582–589.

Danforth, B.N. (1999). Emergence dynamics and bet hedging in a desert bee, *Perdita portalis*. Proceedings of the Royal Society of London. Series B: Biological Sciences, 266(1432), 1985–1994.

Davison, P.J., Field, J. (2018). Limited social plasticity in the socially polymorphic sweat bee *Lasioglossum calceatum*. Behavioral Ecology and Sociobiology, 72, 1–13.

Davies, W.J., Metcalfe, J., Lodge, T.A., da Costa, A.R. (1986). Plant growth substances and the regulation of growth under drought. Functional Plant Biology, 13(1), 105–125.

Fritze, H., Stewart, I.T., Pebesma, E. (2011). Shifts in western North American snowmelt runoff regimes for the recent warm decades. Journal of Hydrometeorology, 12(5), 989–1006.

Frankham, R. (2005). Genetics and extinction. Biological Conservation, 126(2), 131–140.

Frohlich, D.R. (1983). On the nesting biology of *Osmia (Chenosmia) bruneri* (Hymenoptera: Megachilidae). Journal of the Kansas Entomological Society, 56(1), 123–130.

Frohlich, D.R., Tepedino, V.J. (1986). Sex ratio, parental investment, and interparent variability in nesting success in a solitary bee. Evolution, 40(1), 142–151.

Goulson, D., Nicholls, E. (2016). The canary in the coalmine; bee declines as an indicator of environmental health. Science Progress, 99(3), 312–326.

Hung, K.-L.J., Sandoval, S.S., Ascher, J.S., Holway, D.A. (2021). Joint impacts of drought and habitat fragmentation on native bee assemblages in a California biodiversity hotspot. Insects, 12(2), 135.

Jerome, D.K., Petry, W.K., Mooney, K.A., Iler, A.M. (2021). Snow melt timing acts independently and in conjunction with temperature accumulation to drive subalpine plant phenology. Global Change Biology, 27(20), 5054–5069.

Kammerer, M., Goslee, S.C., Douglas, M.R., Tooker, J.F., Grozinger, C.M. (2021). Wild bees as winners and losers: Relative impacts of landscape composition, quality, and climate. Global Change Biology, 27(6), 1250–1265.

Kazenel, M.R., Wright, K.W., Griswold, T., Whitney, K.D., Rudgers, J.A. (2024). Heat and desiccation tolerances predict bee abundance under climate change. Nature, 628(8007), 342–348.

Kudo, G. (1992). Performance and phenology of alpine herbs along a snow-melting gradient. Ecological Research, 7(3), 297–304.

Kuppler, J., Kotowska, M.M. (2021). A meta□analysis of responses in floral traits and flower–visitor interactions to water deficit. Global Change Biology, 27(13), 3095–3108.

LeBuhn, G., Luna, J.V. (2021). Pollinator decline: what do we know about the drivers of solitary bee declines? Current Opinion in Insect Science, 46, 106–111.

McCabe, L.M., Boyle, N.K., Pitts-Singer, T.L. (2024). *Osmia lignaria* (Hymenoptera: Megachilidae) increase pollination of Washington sweet cherry and pear crops. Environmental Entomology, 53(4), 698–705.

Muluneh, M.G. (2021). Impact of climate change on biodiversity and food security: a global perspective—a review article. Agriculture & Food Security, 10(1), 1–25.

Nardini, A. (2022). Hard and tough: the coordination between leaf mechanical resistance and drought tolerance. Flora, 288, 152023.

Newman, D., Pilson, D. (1997). Increased probability of extinction due to decreased genetic effective population size: experimental populations of *Clarkia pulchella*. Evolution, 51(2), 354–362.

Ollerton, J., Winfree, R., Tarrant, S. (2011). How many flowering plants are pollinated by animals? Oikos, 120(3), 321–326.

Outhwaite, C.L., McCann, P., Newbold, T. (2022). Agriculture and climate change are reshaping insect biodiversity worldwide. Nature, 605(7908), 97–102.

Ostwald, M.M., Haney, B.R., Fewell, J.H. (2022). Ecological drivers of non-kin cooperation in the Hymenoptera. Frontiers in Ecology and Evolution, 10, 768392.

Pardee, G.L., Griffin, S.R., Stemkovski, M., Harrison, T., Portman, Z.M., Kazenel, M.R., Lynn, J.S., Inouye, D.W., Irwin, R.E. (2022). Life-history traits predict responses of wild bees to climate variation. Proceedings of the Royal Society B, 289(1973), 20212697.

Phillips, B.B., Shaw, R.F., Holland, M.J., Fry, E.L., Bardgett, R.D., Bullock, J.M., Osborne, J.L. (2018). Drought reduces floral resources for pollinators. Global Change Biology, 24(7), 3226–3235.

Peterson, J.H., Roitberg, B.D., Peterson, J.H. (2006). Impacts of flight distance on sex ratio and resource allocation to offspring in the leafcutter bee, *Megachile rotundata*. Behavioral Ecology and Sociobiology, 59, 589–596.

Peterson, J. H., & Roitberg, B. D. (2006). Impact of resource levels on sex ratio and resource allocation in the solitary bee, Megachile rotundata. Environmental Entomology, 35(5), 1404–1410.

Pitts-Singer, T.L., Artz, D.R., Peterson, S.S., Boyle, N.K., Wardell, G.I. (2018). Examination of a managed pollinator strategy for almond production using *Apis mellifera* (Hymenoptera: Apidae) and *Osmia lignaria* (Hymenoptera: Megachilidae). Environmental Entomology, 47(2), 364–377.

Potts, S.G., Biesmeijer, J.C., Kremen, C., Neumann, P., Schweiger, O., Kunin, W.E. (2010). Global pollinator declines: trends, impacts and drivers. Trends in Ecology & Evolution, 25(6), 345–353.

Potts, S.G., Imperatriz-Fonseca, V., Ngo, H.T., Aizen, M.A., Biesmeijer, J.C., Breeze, T.D., Dicks, L.V. et al. (2016). Safeguarding pollinators and their values to human well-being. Nature, 540(7632), 220–229.

Rering, C.C., Franco, J.G., Yeater, K.M., Mallinger, R.E. (2020). Drought stress alters floral volatiles and reduces floral rewards, pollinator activity, and seed set in a global plant. Ecosphere, 11(9), e03254.

Rose-Person, A., Spasojevic, M.J., Forrester, C., Bowman, W.D., Suding, K.N., Oldfather, M.F., Rafferty, N.E. (2024). Early snowmelt advances flowering phenology and disrupts the drivers of pollinator visitation in an alpine ecosystem. Alpine Botany, 1–10.

Saccheri, I., Kuussaari, M., Kankare, M., Vikman, P., Fortelius, W., Hanski, I. (1998). Inbreeding and extinction in a butterfly metapopulation. Nature, 392(6675), 491–494.

Strzepek, K., Yohe, G., Neumann, J., Boehlert, B. (2010). Characterizing changes in drought risk for the United States from climate change. Environmental Research Letters, 5(4), 044012.

Sun, S., Sun, G., Caldwell, P., McNulty, S., Cohen, E., Xiao, J., Zhang, Y. (2015). Drought impacts on ecosystem functions of the US National Forests and Grasslands: Part II assessment results and management implications. Forest Ecology and Management, 353, 269–279.

Soroye, P., Newbold, T., Kerr, J. (2020). Climate change contributes to widespread declines among bumble bees across continents. Science, 367(6478), 685–688.

Thomson, D.M. (2016). Local bumble bee decline linked to recovery of honey bees, drought effects on floral resources. Ecology Letters, 19(10), 1247–1255.

Vaughan, M., Black, S.H. (2008). Native pollinators: how to protect and enhance habitat for native bees. Native Plants Journal, 9(2), 80–91.

Xu, C., McDowell, N.G., Fisher, R.A., Wei, L., Sevanto, S., Christoffersen, B.O., Weng, E., Middleton, R.S. (2019). Increasing impacts of extreme droughts on vegetation productivity under climate change. Nature Climate Change, 9(12), 948–953.

